# Deubiquitylation and stabilization of p21 by USP11 is critical for cell cycle progression and DNA damage responses

**DOI:** 10.1101/172999

**Authors:** Tanggang Deng, Guobei Yan, Yu Zhou, Xiaoxiao Hu, Jianglin Li, Jun Hu, Hui Zhang, Peifu Feng, Xunan Sheng, Jieying Chen, Hongchang Ma, Yang Sun, Dong Wei, Bin Hu, Jing Liu, Weihong Tan, Mao Ye

## Abstract

p21^WAF1/CIP1^ is a broad-acting cyclin-dependent kinase inhibitor. Its stability is essential for proper cell cycle progression and cell fate decision. Ubiquitylation by the multiple E3 ubiquitin ligases complex is the major regulatory mechanism of p21, which induces p21 degradation. However, it is unclear whether ubiquitylated p21 can be recycled. In this study, we report USP11 as a deubiquitylase of p21. In the nucleus, USP11 binds to p21, catalyzes the removal of polyubiquitin chains conjugated onto p21 and stabilizes p21 protein. As a result, USP11 reverses p21 polyubiquitylation and degradation mediated by SCF^SKP2^, CRL4^CDT2^ and APC/C^CDT20^ in a cell cycle-independent manner. Loss of USP11 causes the destabilization of p21 and induces the G1/S transition in unperturbed cells. Furthermore, p21 accumulation mediated by DNA damage is completely abolished in cells depleted of USP11, which results in abrogation of the G2 checkpoint and induction of apoptosis. Functionally, USP11-mediated stabilization of p21 inhibits cell proliferation and tumorigenesis in vivo. These findings reveal an important mechanism by which p21 can be stabilized by direct deubiquitylation and pinpoint a crucial role of the USP11-p21 axis in regulating cell cycle progression and DNA damage responses.

## Introduction

The cyclin-dependent kinase (CDK) inhibitor p21 (also known as p21^WAF1/Cip1^) is a key negative regulator of cell cycle progression, which mediates cell cycle arrest at the G1 or G2 phase in response to a variety of stress stimuli (1). p21 contributes to the G1 arrest primarily by inhibiting cyclin E and cyclin A/CDK2 activity (2), which results in the hypo-phosphorylation of the retinoblastoma protein (pRb) and inhibits the release and activation of the transcription factor E2F – a protein required for S-phase entry (3). p21 sustains cell cycle arrest at the G2 phase by blocking the interaction between CDK1 and CDK-activating kinase, thus inhibiting the activating phosphorylation of CDK1 at Thr-161 (4). Moreover, several studies have reported that p21 also mediates arrest at G2 by retaining the cyclin B1-CDK1 complex in the nucleus, degrading cyclin B and decreasing the expression of early mitotic inhibitor 1 (Emi1) (5-7).

Under normal growth conditions, p21 is an unstable protein with a relatively short half-life (8, 9). Its degradation is controlled primarily through ubiquitin-proteasome pathway(9). Three E3 ubiquitin ligase complexes SCF^SKP2^, CRL4^CDT2^ and APC/C^CDT20^ have been reported to promote p21 ubiquitylation and degradation in the nucleus. During the G1/S transition, the SCF^SKP2^ complex promotes the ubiquitylation and degradation of p21 after it is phosphorylated at Ser130 by CDK2 (10, 11), whereas the CRL4^CDT2^ complex mediates the ubiquitin-dependent proteolysis of p21 only when p21 is bound to PCNA and phosphorylated at Ser-114 during the S phase (12). When bound to CDK1/cyclin B during prometaphase, p21 is degraded by the APC/C^CDT20^ complex(13). In contrast, p21 stability can be positively regulated by various mechanisms. Phosphorylation of p21 by p38 alpha, JNK1, AKT and NDR has been reported to enhance its stability (14-16). Wisp39, nucleophosmin/B23, hSSB1 and TRIM39 were found to stabilize p21 by engaging in protein-protein interactions (17-20). Cables1 stabilizes p21 by antagonizing PSMA3-mediated proteasomal degradation (21). However, it remains unclear whether ubiquitylated p21 can be recycled.

The removal of ubiquitin from a target protein by deubiquitylase has emerged as an important regulatory mechanism of many cellular functions. The human genome encodes approximately 98 deubiquitylases that can be subdivided into six families (22). USP11 is a deubiquitylase that belongs to the ubiquitin-specific processing protease (USP) family, which is primarily localized to the nucleus and possesses multiple highly conserved domains including Cys box, Asp, KRF and His box (23). Growing evidence has shown that USP11 plays an important role in signal transduction, apoptosis, DNA repair and viral replication by regulating the stability of its substrates (24-27). USP11 dysregulation has been found in a variety of tumors, including colorectal cancer, melanoma, glioma and cervical cancer (28-30).

In this study, we identified USP11 as the first deubiquitylase that directly reverses p21 polyubiquitylation and stabilizes the p21 protein. We also demonstrated that the USP11-p21 axis is critical for regulating cell cycle progression and DNA damage-induced G2 arrest. Our findings reveal an important missing piece regarding the regulation of p21 stability and indicate a previously unknown molecular function of USP11 in controlling cell cycle progression and DNA damage responses.

## Results

### USP11 interacts with p21

USP11 has been shown to function as a deubiquitylating enzyme that stabilizes multiple cellular proteins by cleaving ubiquitin-protein bonds. To search for cellular proteins that interact with USP11, we expressed Flag-tagged USP11 protein in A549 cells and purified USP11-bound protein complexes using an anti-Flag monoclonal antibody coupled to Dynabeads. USP11-associated proteins were identified by liquid chromatography mass spectrometry/mass spectrometry (LC-MS/MS). Intriguingly, p21 was present in the purified USP11 complexes, but not in the control purifications (Fig. S1A). Given the known cellular feature of p21 that can be rapidly degraded by ubiquitylation, we focused our attention on p21 as an interacting protein with USP11. To confirm the interaction between USP11 and p21, Flag-USP11 or Myc-p21 plasmid was transfected into A549 cells, and co-immunoprecipitations (co-IP) was performed using an anti-Flag or anti-Myc antibody. The results showed that p21was detected in the Flag-USP11 immunoprecipitates (Fig. 1A), and that USP11 was present in Myc-p21 immunoprecipitates (Fig. 1B). Meanwhile, the association of endogenously expressed p21 and USP11 was also investigated using co-IP. USP11 and p21 were separately immunoprecipitated from A549 cells, and the reciprocal protein was detected using western blotting. As shown in Figure 1C and D, both USP11 and p21 were detected in their individual immunoprecipitated complexes, but not in the isotype-matched negative control IgG complexes. To determine whether USP11 and p21 directly interact with each other, we generated and purified recombinant USP11 and p21. Purified GST-USP11 but not the GST control was able to bind to GST-p21 under cell-free conditions (Fig. 1E), demonstrating a direct interaction between USP11 and p21. Similar results were obtained by incubating purified GST-USP11 with extracts from A549 cells (Fig. S1B). In addition, the co-localization of both USP11 and p21 occurred in the nucleus (Fig. 1F). Collectively, these results suggest that USP11 physically interacts with p21 *in vivo* and *in vitro*.

**Fig. 1.**
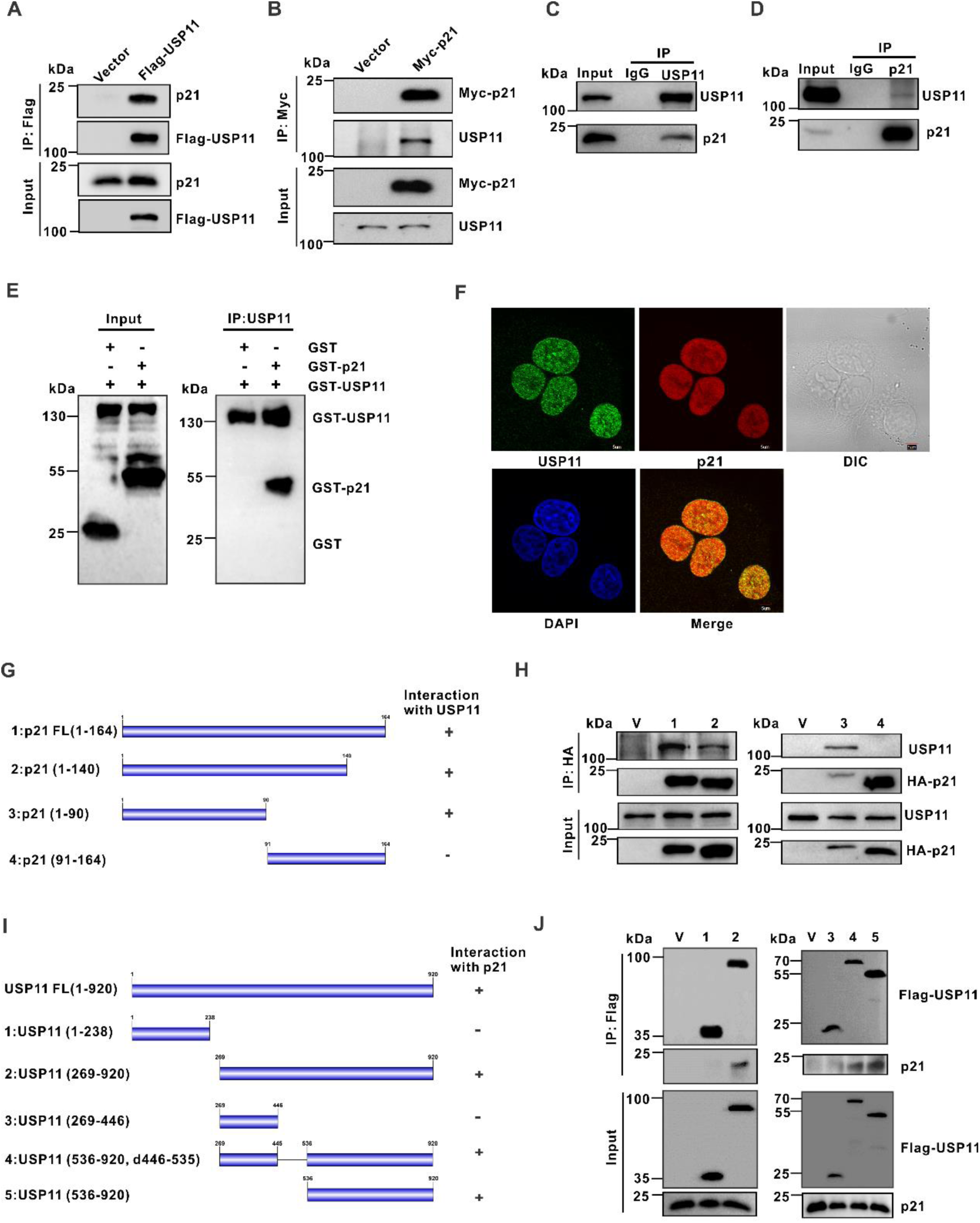
USP11 interacts with p21. (A and B) A549 cells were transfected with plasmids expressing either Flag-USP11 or Myc-p21. Total cell lysates were subjected to immunoprecipitation with anti-Flag (A) or anti-Myc antibody (B). The immunoprecipitates were then probed with the indicated antibodies. (C and D) A549 cell lysates were subjected to immunoprecipitation with control IgG, anti-USP11(C) or anti-p21 (D) antibody. The immunopreciptates were then probed with the indicated antibodies. (E) GST, GST-p21 and GST-USP11 produced from bacteria were assessed using western blotting, and anti-USP11 antibodies were used to immobilize purified GST-USP11 protein onto protein A beads; these beads were then incubated with purified GST or GST-p21. The bound proteins were then eluted and analyzed by western blotting using anti-GST antibodies. (F) The subcellular localization of endogenous USP11 (green) and p21 (red) in A549 cells were visualized using immunofluorescence with anti-USP11 and anti-p21 antibodies. DNA was stained with DAPI, and a merged view of the red and green channels within the same field is shown (merge). (G) Schematic representation of the HA-tagged full-length p21 (FL) and its various deletion mutants. (H) A549 cells transfected with the indicated constructs were lysed. The cell lysates were subjected to immunoprecipitation with anti-HA antibodies, and the immunoprecipitates were then probed with the indicated antibodies. (V: Vector). (I) Schematic representation of the Flag-tagged full-length USP11 (FL) and its various deletion mutants. (J) A549 cells transfected with the indicated constructs were lysed. The cell lysates were subjected to immunoprecipitation with the anti-Flag antibodies, and the immunoprecipitates were then probed with the indicated antibodies. (V: Vector)

To map the USP11-binding region on p21, a series of p21-deletion mutants were expressed in A549 cells (Fig. 1G). Co-IP assays revealed that the N-terminal region (aa 1-90) of p21 was critical for the interaction between USP11 and p21 (Fig. 1H). Conversely, mapping the USP11 region required for p21 binding revealed that the C-terminus (aa 536-920) was responsible for the interaction with p21 (Fig. 1I and J).

### USP11 regulates the protein level of p21

Protein-protein interactions are known to play key roles in regulating p21 levels. Given the identified interaction of USP11 with p21, we next investigated whether USP11 affects the steady-state levels of p21. USP11 was introduced into A549 (p53^+/+^) as well as two HCT116 cell lines with a p53 wild-type (HCT116 WT) and null (HCT116 p53^-/-^) genotype. Interestingly, USP11 overexpression resulted in a significant increase of endogenous p21 levels (Fig. 2A), and increasing USP11 expression caused an elevation of p21 levels in a dose-dependent manner in all cell lines regardless of the p53 status (Fig. 2B and C). In contrast, p53 levels were unaffected by USP11 overexpression, indicating that USP11 increased p21 levels in a p53-independent manner. Notably, overexpression of a catalytically inactive USP11 mutant (C275S/C283S) had no effect on p21 levels (Fig. 2A-C), implying that USP11-mediated upregulation of p21 may depend on the function of USP11 as a deubiquitylaing enzyme. To further confirm the regulation of p21, we performed a loss-of-function analysis using two independent USP11-specific short hairpin RNAs (shRNAs) in the above-mentioned cell lines. As predicted, USP11 knockdown abolished p21 levels without affecting p53 expression (Fig. 3D). Similar results were obtained using USP11-specific small interfering RNAs (siRNAs) in the A549, H460 and HCT116 cell lines (Fig. S2A).

**Fig. 2.**
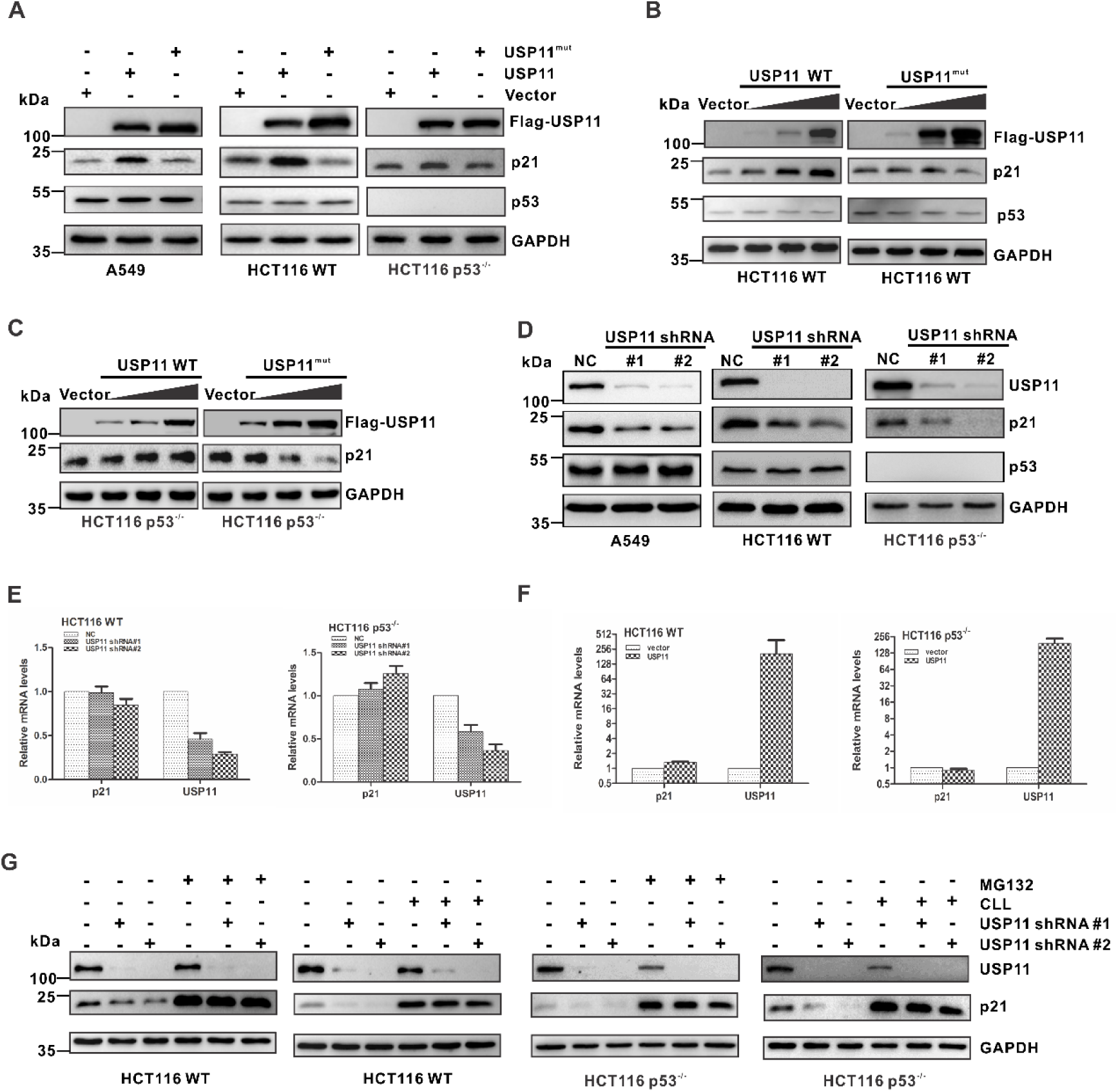
USP11 regulates the protein level of p21. (A) A549, HCT116 WT and HCT116 p53^-/-^ cells were transfected with the indicated constructs. Total protein was extracted and subjected to western blotting using the indicated antibodies. (B and C) Increasing amounts of USP11 WT or USP11^mut^ were transfected into HCT116 WT (B) and HCT116 p53^-/-^ (C) cells, and total protein was extracted from these cells and subjected to western blotting using the indicated antibodies. (D) A549, HCT116 WT and HCT116 p53^-/-^ cells were infected with the indicated lentiviral constructs. The resulting cell extracts were analyzed using western blotting with the indicated antibodies. (E and F) Total RNA from cells either infected with the indicated lentiviral shRNAs (E) or cells transfected with the indicated constructs (F) was isolated and subjected to qRT-PCR. The error bars represent the standard deviation (SD) of triplicate measurements. (G) HCT116 WT and HCT116 p53-/- cells infected with the indicated lentiviral shRNAs were treated with DMSO, MG132 (20 μM) or CLL (Clasto-Lactacystin β-lactone, 10 μM) for 6 h, and the indicated proteins were analyzed using western blotting.

The effect of USP11 on the p21 steady-state levels was not due to changes in transcription because neither USP11 knockdown nor overexpression affected the p21 mRNA levels (Fig. 3E and F; Fig. S2B), indicating that USP11 does not regulate p21 expression at the transcriptional level. Furthermore, downregulation of p21 caused by USP11 knockdown could be blocked by the proteasome inhibitor MG132 and CLL (Fig. 3G and Fig. S2C), suggesting that USP11 maintains the steady-state levels of p21 by blocking its proteasomal degradation.

**Fig. 3.**
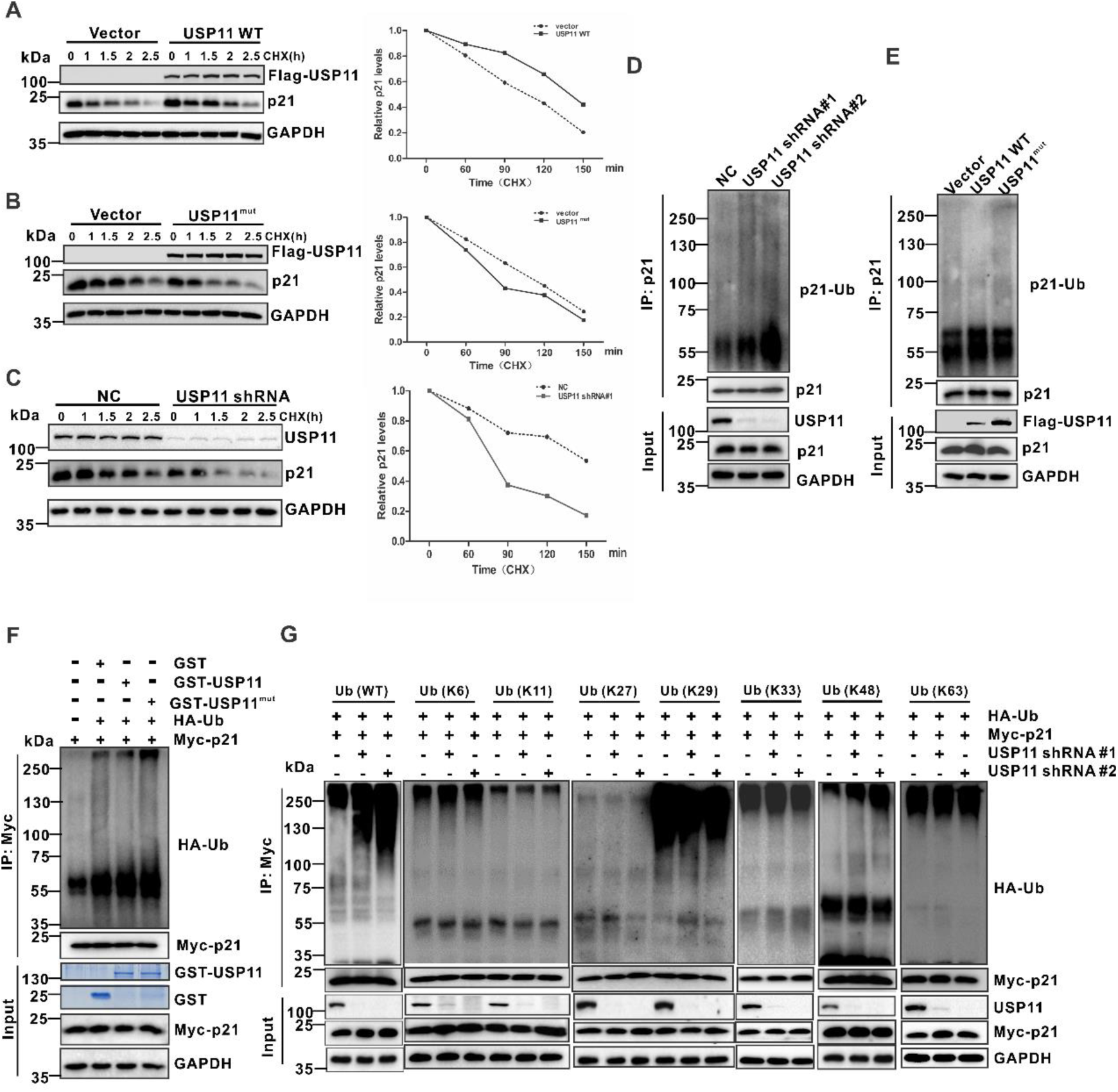
USP11 stabilizes p21 via deubiquitylation. (A and B) HCT116 WT cells transfected with the indicated constructs were treated with 50 μg ml^-1^ cycloheximide (CHX), collected at the indicated time points and immunoblotted with the indicated antibodies. Quantification of the p21 levels relative to GAPDH expression is shown. (C) HCT116 WT cells infected with the indicated lentiviral shRNAs, were treated with 50 μg ml^-1^ CHX, collected at different time points and then immunoblotted with the indicated antibodies. Quantification of the p21 levels relative to GAPDH expression is shown. (D and E) HCT116 WT cells either infected with the indicated lentiviral shRNAs (D) or transfected with the indicated constructs (E) were treated with MG132 (20 μM) for 6 h prior to harvest. p21 was immunoprecipitated with an anti-p21 antibody, and the immunoprecipitates were probed with the indicated antibodies. (F) Ubiquitylated Myc-p21 was incubated with either GST-tagged USP11 WT or the GST-tagged USP11^mut^ purified from bacteria using glutathione agarose. After co-incubation, Myc-p21 was immunoprecipitated using an anti-Myc antibody, and the immunoprecipitates were probed using antibodies against HA and Myc. Recombinant GST-USP11 was purified from bacteria and analyzed using SDS-PAGE and Coomassie blue staining. (G) Myc-p21 and various HA-ubiquitin mutants were transfected into HCT116 cells infected with the indicated lentiviral shRNA for 24h. The cells were treated with 20 μM the proteasome inhibitor MG132 (Sigma) for 6 h. Myc-p21 was immunoprecipitated with an anti-myc antibody, and the immunoprecipitates were probed with the indicated antibodies.

### USP11 stabilizes p21 by deubiquitylation

Because USP11 regulates the protein levels of p21, we questioned whether USP11 stabilizes p21. To this end, in the presence or absence of Flag-USP11, cells were treated with cycloheximide (CHX) to inhibit protein biosynthesis, and protein extracts obtained at indicated time points were analyzed. We found that overexpression of wild-type USP11but not catalytically inactive mutant profoundly extended the half-life of the p21 protein (Fig. 3A and B; Fig. S3A). Conversely, knockdown of USP11 resulted in a significant decrease in the half-life of p21 (Fig. 3C and Fig. S3B). To further understand the underlying mechanism whereby USP11 regulates the stability of p21, we measured the levels of polyubiquitylation of p21 in HCT116 cells. Silencing USP11 expression using two independent shRNAs led to a significant increase in p21 polyubiquitylation (Fig. 3D), whereas the overexpression of wild-type USP11 reduced the levels of polyubiquitylated p21 (Fig. 3E). In contrast, the catalytically inactive mutant failed to protect p21 from ubiquitylation (Fig. 3E), suggesting that the enzymatic activity of USP11 is essential for the USP11-dependent deubiquitylation of p21. To verify that p21 is a direct substrate of USP11, we purified USP11 and ubiquitylated p21, and incubated these two proteins in a cell-free system. As expected, wild-type USP11 but not the catalytically inactive mutant decreased p21 polyubiquitylation in vitro (Fig. 3F). These data indicate that USP11 directly deubiquitylates p21.

To investigate the type of poly-Ub chain on p21 that is removed by USP11, we transfected HCT116 cells with Myc-tagged p21, together with HA-tagged ubiquitin mutants in which all lysines, except only one lysine (K6, K11, K27, K29, K33, K48 or k63), were mutated into arginines. As shown in Figure 3G, USP11 knockdown significantly increased K48-linked poly-Ub but not any other isopeptide-linked (K6, K11, K27, K29, K33 or K63) poly-Ub. The result suggests that USP11 removes K48-linked poly-Ub in p21.

### USP11 stabilizes p21 in response to DNA damage

p21 can be induced under DNA damage condition via p53-dependent and p53-independent pathways. To explore whether USP11 is involved in the DNA damage-mediated regulation of p21, we treated cells with genotoxic agents. In agreement with previous reports, etoposide treatment led to the upregulation of p21 levels in HCT116 WT and HCT116 p53^-/-^ cells (Fig. 4A and B). Intriguingly, etoposide-induced p21 accumulation was significantly abolished in USP11-depleted cells (Fig. 4A and B). Similarly, USP11 knockdown also significantly decreased the p21 elevation triggered by doxorubicin (Fig. 4A and B). Notably, depletion of USP11 did not abolish the induction of p21 mRNA in response to genotoxic treatment (Fig. 4C and D). Collectively, these findings suggest that USP11 is indispensable for the expression of p21 under physiological conditions as well as in response to DNA damage (Fig. 4E).

**Fig. 4.**
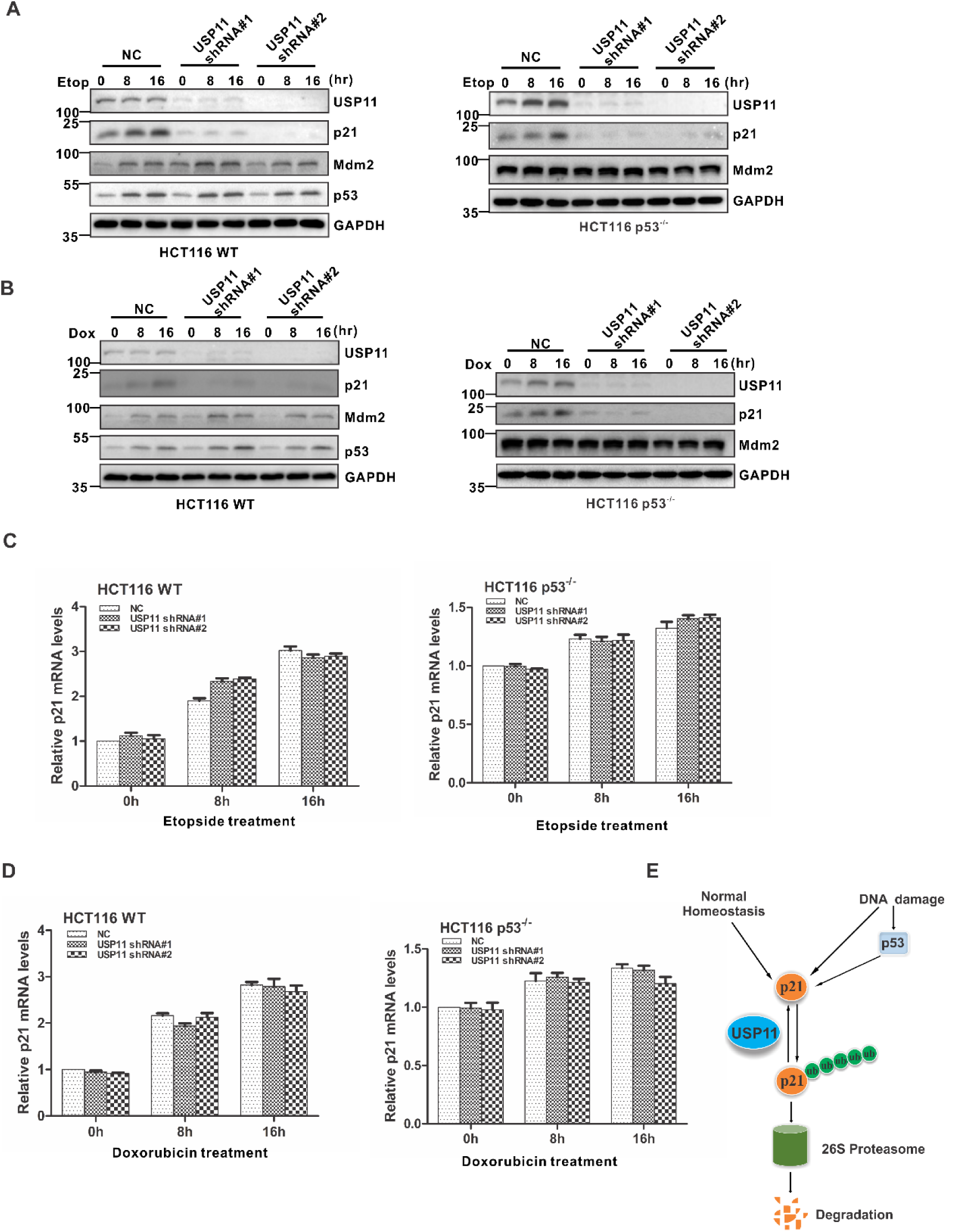
USP11 knockdown abolishes p21 elevation triggered by genotoxic agents. (A and B) HCT116 WT (A) and HCT116 p53^-/-^ (B) cells infected with the indicated lentiviral shRNAs were treated with 5 μM etoposide (Etop) or 0.2 μM doxorubicin (Dox) for either 8 or 16 h. Cell lysates were then extracted and subjected to western blotting. (C and D) Total RNA from HCT116 WT (C) and HCT116 p53^-/-^ (D) cells infected with the indicated lentiviral shRNAs and treated with 5 μM etoposide (Etop) or 0.2 μM doxorubicin (Dox) for either 8 or 16 h was isolated and subjected to qRT-PCR. The error bars represent the SD of triplicate measurements. (E) A proposed working model for p21 regulation by USP11 in response to DNA damage.

### USP11 protects p21 from ubiquitin-mediated degradation in a cell cycle-independent manner

Three E3 ubiquitin ligase complexes SCF^SKP2^, CRL4^CDT2^ and APC/C^CDC20^ have been reported to induce p21 ubiquitylation and degradation at different phases during an unperturbed cell cycle. To assess which E3 ubiquitin ligase complex is regulated by USP11, HCT116 cells stably expressing the indicated shRNAs were synchronized at each phase (Fig. S4). Strikingly, USP11 knockdown led to a significant decrease of p21 at all phases of the cell cycle, although the p21 protein level varied during the cell cycle (Fig. 5A). Furthermore, we examined whether the effect of USP11 on p21 was associated with SCF^SKP2^, CRL4^CDT2^ or APC/C^CDC20^. Knockdown of USP11 using shRNAs significantly decreased p21 levels with concomitant increases in SKP2 (Fig. 5B), but the levels of CDT2 and CDC20 were unchanged (Fig. 5C and D). However, when SKP2, CDT2 or CDC20 was knocked down by siRNA, USP11 depletion-induced p21 degradation and ubiquitylation was abolished (Fig. 5B-G). Altogether, these results indicate that USP11 stabilizes p21 via the reversal of SCF^SKP2^, CRL4^CDT2^ or APC/C^CDC20^-mediated ubiquitylation and degradation in a cell cycle-independent manner (Fig. 5H).

**Fig. 5.**
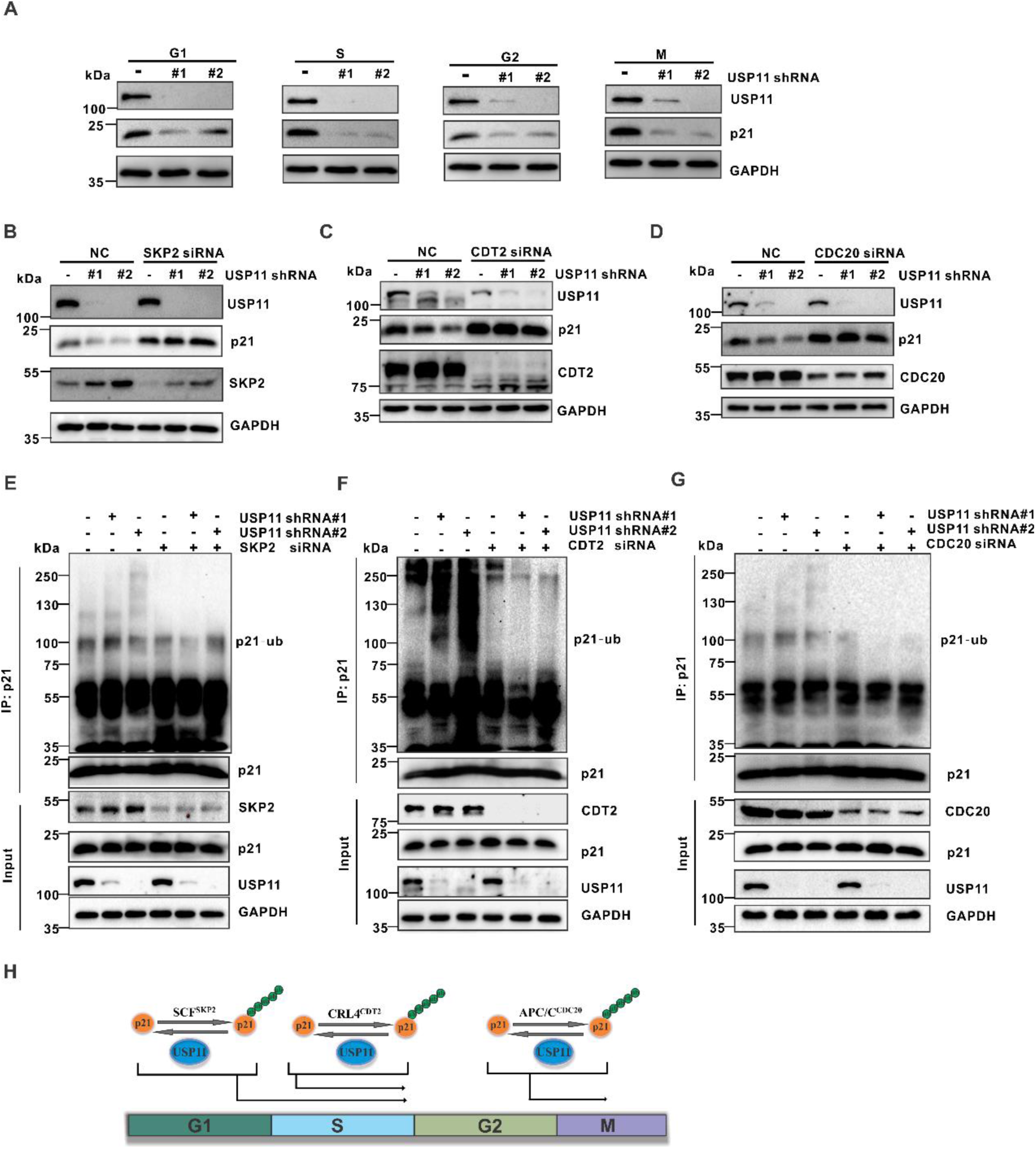
USP11 protects p21 from ubiquitin-mediated degradation. (A) HCT116 WT cells infected with the indicated lentiviral shRNAs were synchronized using a double thymidine block and released at the indicated phases. The resulting cell lysates were subjected to western blotting using the indicated antibodies. (B-D) HCT116 WT cells infected with the indicated shRNAs were transfected with scrambled, SKP2 (B), CDT2 (C), or CDC20 (D) siRNA for 48h, after which the cell lysates were harvested and analyzed using western blotting. (E-G) HCT116 cells infected with the indicated shRNAs were transfected with scrambled, SKP2 (E), CDT2 (F) and CDC20 (G) siRNA as indicated for 48h and treated with 20 μM the proteasome inhibitor MG132 (Sigma) for another 6 h. p21 was immunoprecipitated with an anti-p21 antibody, and the immunoprecipitates were probed with the indicated antibodies. (H) A proposed working model that illustrates how USP11 reverse p21 ubiquitination in a cell-cycle independent manner.

### USP11 regulates cell cycle progression and the DNA damage response in a p21-dependent manner

Because p21 regulates cell cycle progression at G1 phase, we hypothesized that USP11 may affect cell cycle progression from G1 to S phase. To test this hypothesis, the percentage of cells in S phase was determined by measuring the DNA content and incorporation of BrdU, as well as by performing double thymidine block and release. As predicted, the percentage of cells in S phase was increased when USP11 was knocked down in HCT WT and HCT116 p53^-/-^ cells (Fig. 6A-C; Fig. S5). In contrast, USP11 depletion in HCT116 p21^-/-^ cells exhibited no effects on the G1/S transition (Fig. 6A and B; Fig. S5), but USP11-depleted cells transfected with exogenous p21 fully prevented the G1/S transition induced by USP11 ablation (Fig. 6D). These results strongly suggest that the USP11-mediated G1/S transition is dependent on p21.

**Fig. 6.**
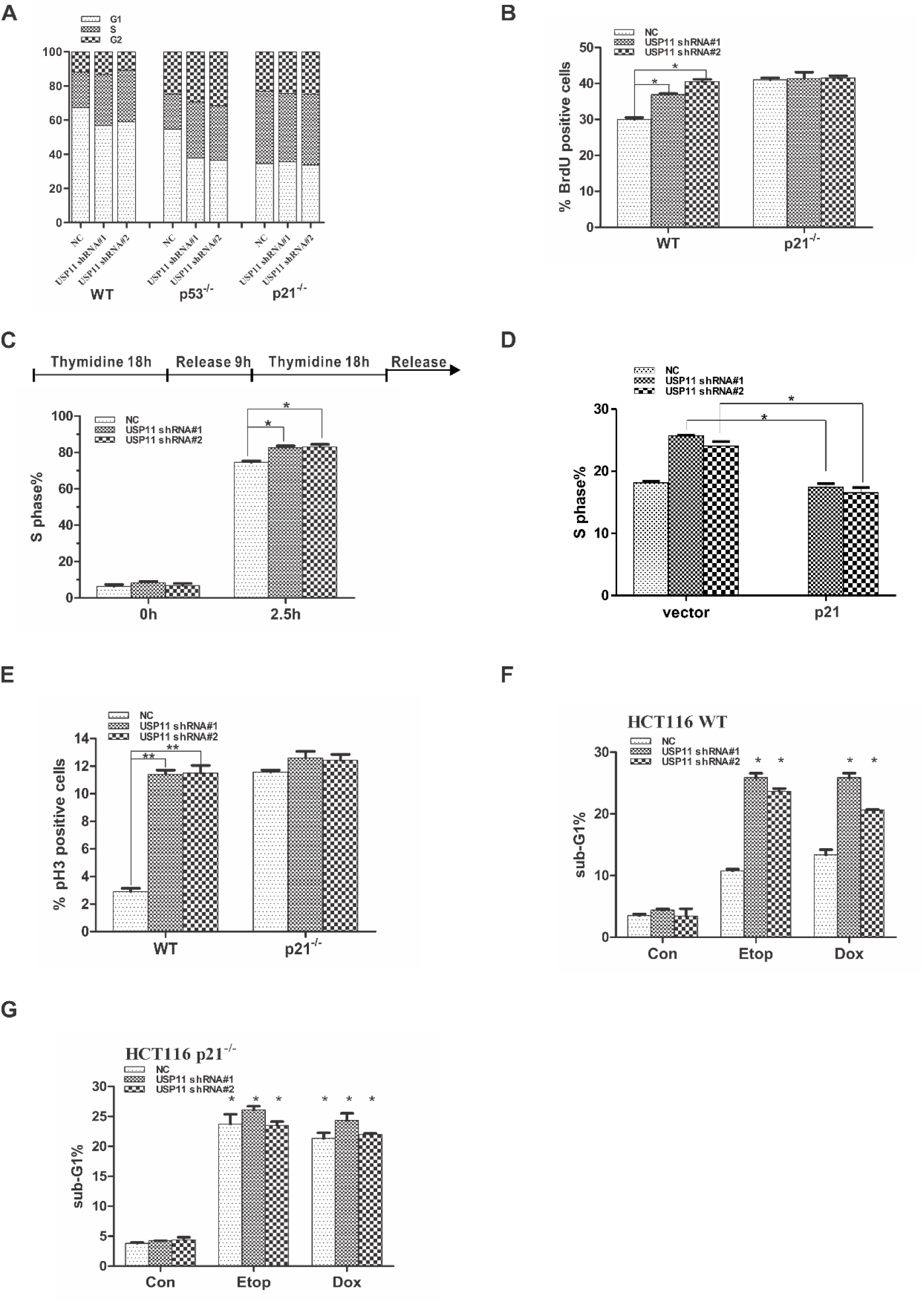
USP11 regulates the G1/S transition and DNA damage-induced G2 checkpoint in a p21-dependent manner. (A) HCT116 WT, HCT116 p53^-/-^ and HCT116 p21^-/-^ cells infected with the indicated lentiviral shRNAs were stained with propidium iodide and analyzed using flow cytometry. (B) HCT116 WT and HCT116 p21^-/-^ cells transfected with the indicated shRNAs were labeled with BrdU for 60 min before harvesting and then analyzed using flow cytometry. The error bars represent the mean ± SD of three independent experiments. *P <0.05. (C) HCT116 WT cells infected with the indicated lentiviral shRNAs were synchronized using a double thymidine block and release. The released cells were then harvested at the indicated times and analyzed using flow cytometry. The error bars represent the mean ± SD of three independent experiments. *P <0.05. (D) HCT116 WT cells infected with the indicated lentiviral shRNAs were transfected with the indicated constructs for 24 h. Cells were stained with propidium iodide and analyzed using flow cytometry. The error bars represent the mean ± SD of three independent experiments. *P <0.05. (E) HCT116 WT and HCT116 p21^-/-^ cells infected the indicated lentiviral shRNAs were pretreated with 0.2 μM doxorubicin for 2 h, followed by synchronization with nocodazole (100 ng mL^-1^) for 16 h. The mitotic index was determined using pH3 staining as a marker of mitosis. The error bars represent the mean±SD of three independent experiments. **P < 0.01. (F and G) HCT116 WT (F) and HCT116 p21^-/-^ (G) cells were infected with the indicated lentiviral shRNAs, followed by treatment with either 0.2 μM doxorubicin (Dox) or 5 μM etoposide (Etop) for 48 h, and subsequent flow cytometry analysis of the sub-G1 fraction. The error bars indicate the mean±SD of three independent experiments. *P < 0.05.

To determine whether p21 is required for the function of USP11 in the G2/M checkpoint after DNA damage, cells were treated with a low-dose of doxorubicin. The phopho-histone3 (pH3) at Ser10, an indicator of cells at M phase, was used to monitor the G2/M checkpoint. As shown in Figure 6E and Figure S6, after doxorubicin treatment, the percentage of cells in M phases was significantly increased in HCT WT cells with USP11 knockdown. However, in HCT116 cells lacking p21 (HCT116 p21^-/-^), silencing USP11 had no effect on the increased percentage of cells in M phase, indicating that USP11 depends on p21 to sustain the DNA damage-induced G2/M checkpoint.

To investigate the effect of USP11 on apoptosis induced by a DNA-damaging agent, HCT116 WT and HCT116 p21^-/-^ cells were treated with either doxorubicin or etoposide. The percentage of cells in sub-G1 phase (apoptotic cells) was measured using flow cytometry with propidium iodide staining. Compared with the control cells, USP11-depleted HCT116 WT cells exhibited a significant increase in the levels of apoptosis after a 24-h treatment with either doxorubicin or etoposide (Fig. 6F and G). Interestingly, USP11 knockdown did not affect apoptosis triggered by either doxorubicin or etoposide in cells lacking p21 (Fig. 6F and G). Collectively, these data show that USP11 knockdown sensitizes cells to DNA damage-induced apoptosis by abolishing p21 accumulation.

### Loss of USP11 promotes tumor cell growth via the downregulation of p21

To investigate whether USP11 functions as a tumor suppressor by regulating p21, USP11-depleted A549 cells were implanted into nude mice and tumor growth was monitored at the indicated time points. Compared with mice implanted with control shRNA-infected cells, mice bearing USP11-shRNA expressing A549 cells showed increased tumor growth throughout the experiment (Fig. 7A). At 37 days after tumor cell implantation, the volume and weight of the tumor formed by USP11-depleted A549 cells were significantly increased. (Fig. 7A-C). The results of an immunohistochemical analysis verified the reduced expression of USP11 and p21 in the xenograft tumors (Fig. 7D). Next, we analyzed the effect of USP11 on cell proliferation. We found that USP11 depletion promoted the proliferation of A549 and HCT116 WT cells, and that p21 restoration completely reversed the effect of USP11 depletion (Fig. 7E). Notably, USP11 knockdown showed no effect on the proliferation of HCT116 p21^-/-^ cells (Fig.7E). Conversely, overexpression of USP11, but not the catalytically inactive mutant of USP11, inhibited the proliferation of A549 and HCT116 WT cells (Fig. 7F). Similarly, overexpression of USP11 had no effect on the proliferation of HCT116 p21^-/-^ cells (Fig. 7F). Therefore, the loss of USP11 results in increased proliferation of tumor cells via the downregulation of p21.

**Fig. 7.**
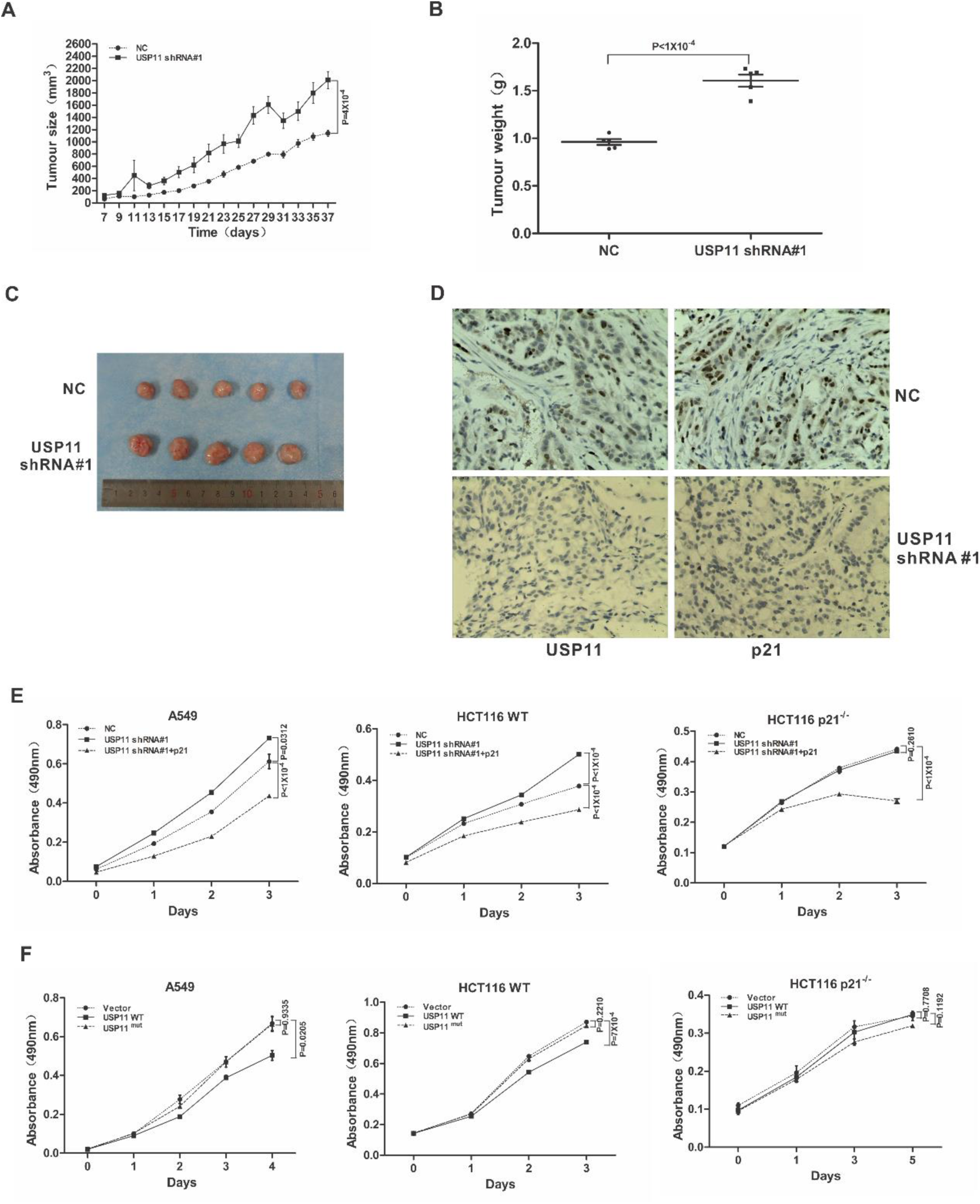
Loss of USP11 promotes tumor growth through the downregulation of p21. (A-C) Tumor growth of 5×10^6^ A549 cells transfected with the indicated shRNAs and subcutaneously injected into mice. Tumor growth (A), tumor weight (B) and tumor images (C) are shown. (D) Immunohistochemical analysis of USP11 and p21 expression in tumor xenografts. Formalin-fixed sections of tumors formed 37 days after injection of A549 cells infected with the indicated lentiviral shRNAs were stained with anti-USP11 and anti-p21 antibodies. Typical fields of view are presented. (E) A549, HCT116 WT and HCT116 p21^-/-^ cells were infected with the indicated lentiviral shRNAs, and then transfected with the indicated constructs. Cell proliferation was monitored using MTT assays at the indicated time points. Statistical significance was determined by a two-tailed, unpaired Student’s t-test. (F) A549, HCT116 WT and HCT116 p21^-/-^ cells were transfected with the indicated constructs, and cell proliferation was monitored using MTT assays at the indicated time points.

## Discussion

In the present study, we identified USP11 as a p21 deubiquitylase. Our results indicate that USP11 and p21 interact with each other and colocalize in the nucleus. Overexpression of USP11 stabilizes p21 by removing its ubiquitin chain, whereas USP11 downregulation decreases p21 levels, which is accompanied by increasing ubiquitylation. To the best of our knowledge, this is the first evidence that p21 can be stabilized by direct deubiquitylation mediated by a deubiquitylase.

p21 is a well-known transcriptional target of p53. In response to various stresses including DNA damage and oxidative stress, activation of p53 induces p21 protein expression by binding its promoter (31). A recent study revealed that USP11 deubiquitylates and stabilizes p53 (32). However, our results indicated that USP11 had no effect on p53. USP11 overexpression or knockdown of USP11 failed to affect p53 levels, which is consistent with a previous study demonstrating that USP11 does not interact with p53 and exhibit any effect on the levels of p53 ubiquitylation or stabilizing p53 (33). Furthermore, we found USP11 exerts its function on p21 both in p53 wild-type and null cell lines, suggesting that USP11 regulates p21 levels in a p53-independent manner. In response to genotoxic treatment, p21 was accumulated in p53 wild-type and null cell lines. Strikingly, USP11 knockdown completely abolished p21 elevation induced by genotoxic agents, but not p21 mRNA induction. This finding reveals the interesting fact that the stability of p21 mediated by USP11 is indispensable for both p53-dependent and p53-independent transactivation of p21.

p21 is an unstable protein with a relatively short half-life that can respond to rapid intrinsic and extrinsic alterations (9). Its stability is regulated mainly by post-translational modifications such as phosphorylation and ubiquitylation (34). For the ubiquitin-dependent pathway, three E3 ubiquitin ligase complex, SCF^SKP2^, CRL4^CDT2^ and APC/C^CDT20^, have been identified to promote p21 ubiquitylation and degradation at specific stages of the cell cycle. SCF^SKP2^ is necessary for p21 degradation at the G1/S transition as well as during S phase of the cell cycle (35), whereas CRL4^CDT2^ specifically targets p21 for degradation in S phase (36). During mitosis, the APC/C^CDT20^ complex primarily drives p21 degradation (13). Here, we showed that USP11 protected p21 from ubiquitin-mediated degradation by abolishing the action of the above E3 ubiquitin ligase complex. Loss of p21 expression upon USP11 knockdown was significantly ameliorated by depleting SKP2, CDT2 and CDC20, indicating that USP11-mediated protection of p21 is independent of the cell cycle. Notably, knocking down USP11 caused a significant increase of SKP2 levels without affecting either CDT2 or CDC20 expression. Given that the change in SKP2 levels coincides with the functional activity of USP11 with regard to p21 stability, we speculated that SKP2 may contribute to a positive feedback regulation that enhances the effects of USP11 on p21, but the detailed mechanism underlying USP11-dependent regulation of SKP2 remains to be elucidated.

p21 plays a critical role in cell cycle progression and the cellular response to DNA damage by arresting cell cycle progression at the G1/S and G2/M transitions. Consistent with this notion, our results showed that USP11 depletion promoted the G1/S transition in unperturbed cells, which could be reversed by the ectopic expression of p21. Similar results were obtained in HCT116 p53^-/-^ cells, which is consistent with the result that USP11 regulates the protein levels of p21 in a p53-independent manner. In response to DNA damage, USP11 knockdown abrogates the G2/M checkpoint in cells exposed to genotoxic treatment, which promotes mitotic entry and induces apoptosis. Notably, USP11 knockdown in HCT116 p21^-/-^ cells had no effect on the G1/S transition or DNA damage-induced G2/M checkpoint, suggesting that USP11 is involved in regulating the cell cycle through p21. Collectively, these findings indicated that the USP11-p21 axis plays an important role in regulating cell cycle progression and DNA damage-induced G2 arrest.

It has been reported that p21 can function as a tumor suppressor as well as an oncogene. This dual behavior of p21 primarily depends on its subcellular location. The tumor-suppressive activities of p21 are associated with its nuclear localization, whereas cytoplasmic p21 contributes to its oncogenic effects (9, 37, 38). Our results show that USP11 acts as tumor suppressor, as overexpression of USP11 inhibited cell proliferation, whereas cells with USP11 depletion exhibited increased proliferation. This is consistent with previous reports that USP11 functions as a tumor suppressor (30, 39, 40). Furthermore, silencing USP11 in HCT116 p21^-/-^ cells had no effect on the proliferation, indicating that USP11 exerts its function via p21. Given that USP11 interacts with p21 in the nucleus, we speculated that the biological function of USP11 is associated with the tumor-suppressive activities of nuclear p21. Further studies are necessary to establish a detailed association between USP11 and a variety of human cancers, which will provide clue as to how to utilize USP11 as a potential cancer therapeutic target.

## Materials and Methods

### Cell culture and reagents

HEK293 cells were cultured in DMEM (GIBCO cat. 8116490) supplemented with 10% (vol/vol) FBS; A549 cells were cultured in RPMI 1640 (GIBCO cat. 8116491) supplemented with 10% FBS and HCT116 WT, HCT116 p53^-/-^, and HCT116 p21^-/-^ cells were cultured in McCoy’s 5A medium (Invitrogen cat. 16600082) supplemented with 10% FBS. The following reagents were used: MG132 (Sigma cat. C2211-5MG), nocodazole (Sigma cat. M1404-10MG), GSH-Sepharose (Thermo Fisher Scientific cat. 16100), doxorubicin (Sigma cat. D1515-10MG), and etoposide (Sigma cat. E1383-25MG).

### Immunofluorescence staining

Cells were cultured in Lab-Tek chambers for 24 h, washed three times with phosphate-buffered saline (PBS), and fixed with 4% paraformaldehyde for 15 min. The fixed cells were then washed twice with PBS, permeabilized for 10 min with 0.2% Triton X-100, blocked for 1 h in 5% BSA and incubated with the appropriate primary antibodies overnight at 4°C. The USP11 (1:200) and p21 (1:200) antibodies were used to detect endogenous protein expression. Cells were labeled for 1 h using secondary antibodies conjugated to either FITC (Thermo Fisher Scientific cat. 20-58-060111) or Texas Red (Thermo Fisher Scientific cat. 19-183-062312) at room temperature followed by three washes with PBS. Cells were then counterstained with DAPI at room temperature for 5 min to visualize nuclear DNA. Fluorescence images were acquired using a confocal microscope (Zeiss LSM510).

### Western blotting and immunoprecipitation

Western blotting and co-IP were performed as described previously. Briefly, cells were lysed in M-PER buffer (Thermo Fisher Scientific cat. 78501) containing protease inhibitors (Biotool cat. B14001), and the clarified lysates were resolved on 12% gels using SDS-PAGE and transferred to nitrocellulose membranes for western blotting using ECL detection reagents (Advansta cat. 160625-66). Alternatively, 3 mg of clarified lysates was incubated overnight at 4°C with 3 μg of either relevant primary antibodies or an isotype-matched negative control IgG. Subsequently, the samples were incubated for 1 h with 30 μl of magnetic beads conjugated with protein G (Invitrogen cat.10004D) and then washed four times with co-IP/wash buffer. Precipitated proteins were dissolved in 2× SDS sample buffer, boiled, and subjected to western blot analysis.

### GST pulldown assays

Bacterially expressed GST-USP11 was retained on glutathione sepharose beads (Promega cat. V8611) and incubated for 1 h at 4°C with A549 cells extracts.

### Protein half-life assays

Cells were treated with cycloheximide (50 μg ml^-1^) for various periods of time to block protein synthesis. Crude extracts were prepared, and the protein levels were assessed using western blot analysis.

### Synchronization of HCT116 WT cells

To synchronize HCT116 WT cells at the G1/S border, cells were treated with 2 mM thymidine (Millipore cat. 6060-5mg) for 18 h. Cells were released from the block by washing with PBS followed by the addition of complete growth medium. After 9 h, thymidine was added to the medium to a final concentration of 2 mM, and the cells were cultured for an additional 18 h. Cells were then rinsed twice with PBS, cultured in complete growth medium for 3 h (S-phase cells), 6 h (G2-phase cells) or treated with culture media containing 100 ng ml^-1^ of nocodazole for 11 h (M-phase cells). Cells were collected and analyzed using flow cytometry and western blotting.

## Acknowledgments

We would like to thank Han You for generously providing the HCT116 p21^-/-^ and HCT116 p53^-/-^ cells. This work was supported by grants from the National Basic Research Program of China (No. 2013CB932702), the National Natural Science Foundation of China (Nos. 81171950, 81272220, 81402304 and 81672760), the Interdisciplinary Research Program of Hunan University and the Program for New Century Excellent Talents in University (NCET-13-0195).

### Competing financial interests

The authors declare no competing financial interests exist.

